# An exon-capture system for the entire class Ophiuroidea

**DOI:** 10.1101/014613

**Authors:** Andrew F. Hugall, Timothy D. O’Hara, Sumitha Hunjan, Roger Nilsen, Adnan Moussalli

## Abstract

We present an exon-capture system for an entire class of marine invertebrates, the Ophiuroidea, built upon a phylogenetically diverse transcriptome foundation. The system captures ~90 percent of the 1552 exon target, across all major lineages of the quarter-billion year old extant crown group. Key features of our system are: 1) basing the target on an alignment of orthologous genes determined from 52 transcriptomes spanning the phylogenetic diversity and trimmed to remove anything difficult to capture, map or align, 2) use of multiple artificial representatives based on ancestral states rather than exemplars to improve capture and mapping of the target, 3) mapping reads to a multi-reference alignment, and 4) using patterns of site polymorphism to distinguish among paralogy, polyploidy, allelic differences and sample contamination. The resulting data gives a well-resolved tree (currently standing at 417 samples, 275,352 bp, 91% data-complete) that will transform our understanding of ophiuroid evolution and biogeography.

## INTRODUCTION

Next-generation sequencing is revolutionising phylogenetics through the provision of massive amounts of sequence data (Lemmon and Lemmon 2013). Many studies use transcriptomes to focus sequence effort on a common set of phylogenetically useful markers but RNA sequencing has demanding requirements on sample quality, effectively ruling out many taxa. On the other hand, museums possess many samples suitable for DNA extraction (e.g. in >70% ethanol). To expedite the construction of tree of life phylogenies or to explore the origins and biogeography of biota from remote environments, such as the deep sea, this reservoir of material is vital.

Hybridisation enrichment has emerged as a key technology for collecting targeted DNA sequences from museum specimens (Lemmon and Lemmon 2013). Extracted DNA is fragmented, ligated with adaptors and barcodes, hybridised to probes (or ‘baits’) to enrich or ‘capture’ the targeted sequences, which are then sequenced using next-generation technology (Lemmon and Lemmon 2013). The key limitation is that hybridisation capture is most effective if the genetic distance between probe and target is less than approximately 12% (Hancock-Hanser et al. 2013; although see Li et al. 2013). Thus probes have to be designed from known genetic sequences that are not expected to be too divergent from the target.

For large tree of life-scale phylogenies this hybridisation limitation raises the question of how to capture recognizably orthologous targets across a wide range of phylogenetic divergences. Various approaches have been taken to deal with these issues (see Lemmon and Lemmon 2013). One approach has been to use highly conserved sequences as ‘anchors’ that allow capture across a wide range of taxa, relying upon associated variable flanking regions to provide most of the phylogenetic information (e.g. Ultra-Conserved-Elements: Siepel et al. 2005; Faircloth et al. 2012; Anchored Elements: Lemmon et al. 2012). This deals with the capture problem but requires comparative genome information to establish the candidate set of ‘anchors’ and *post-hoc* ascertainment of flanking region orthology. Another approach is to use known common genes, which directly provide the phylogenetic information but may be too variable to capture over more than a moderate phylogenetic range (Bi et al. 2012; Hedtke et al. 2013; Mandel et al. 2014). Certainly at the class or order level, multiple versions of probes will be required to capture the phylogenetic diversity (Lemmon and Lemmon 2013). Fortunately, for taxonomic groups without genomic-scale data, this diversity of probes can be designed directly from transcriptome-based phylogenetic datasets, albeit restricted to a more limited set of commonly expressed exons. Here we embrace this approach for the marine invertebrate class Ophiuroidea, using probes based on multiple clade-based artificial representatives.

Abundant in marine benthic habitats, ophiuroids (brittlestars, basketstars) are a key group for the study of marine biogeography and macroecology (O’Hara 2007; O’Hara et al. 2011; Stöhr et al. 2012; O’Hara et al. 2014a), especially of the deep sea. However, existing molecular data has been very limited in taxonomic and genetic scope, consisting predominantly of short sequences of mitochondrial (COI, 16S) and/or ribosomal (28S, 18S) DNA. There is no sequenced ophiuroid genome and the closest available (the sea urchin *Stronglyocentrotus purpuratus)* diverged at least 485 mya (Sprinkle and Guensburg 2004). Consequently, we embarked on a three-stage plan to generate a tree of life for the Ophiuroidea, founded on transcriptome data of 425 protein coding genes from 52 taxa across all major groups (O’Hara et al. 2014b), followed by exon-capture developed from this transcriptome gene set. A tapestry approach can then be used to add species with only legacy mtDNA sequences to the tree.

This paper reports on the second stage of the process, the construction and function of probe kits designed to capture and reconstruct variable protein-coding exon data from hundreds of samples across an entire class of marine invertebrates. In effect we combined and extended elements of several existing approaches. We designed our probes from cheaper transcriptome rather than genomic data (Bi et al. 2012) and, rather than searching for conserved elements, we used multiple versions of each probe (Lemmon et al. 2012) in an attempt to limit the genetic distance between any probe and target to less than 12%. We also targeted the mitochondrial COI gene as well as nuclear exons (Hancock-Hanser et al. 2013) to help verify sample identity (by matching against available ‘barcodes’) and to allow incorporation of taxa sequenced with legacy COI data.

## RESULTS

### Probe design

We identified exons from a 425 aligned gene dataset (O’Hara et al. 2014b) derived from 52 ophiuroid transcriptomes and outgroups and used the closest genome *(Strongylocentrotus purpuratus)* as the basis for breaking up the ophiuroid transcriptome data into putative exons. After excluding exons with insufficient sequence length (<99 bp), excessive length variation, repeat elements, or missing data, our final target consisted of 1552 nominal exons in 418 genes spanning 285,165 sites (fig. 1 “final”). The 1552 exons contained 139 thousand variable sites, two-thirds of which were 3^rd^ position. There was a mix of conserved and variable exons, with half having >17% differences across the class. We also included the mitochondrial COI gene (also derived from the transcriptomes). All selected exons could be distinguishable from one another (greater than 24% different) at the level of short read lengths of 100-150 bases (i.e. should not confound one another).

**Fig. 1.**
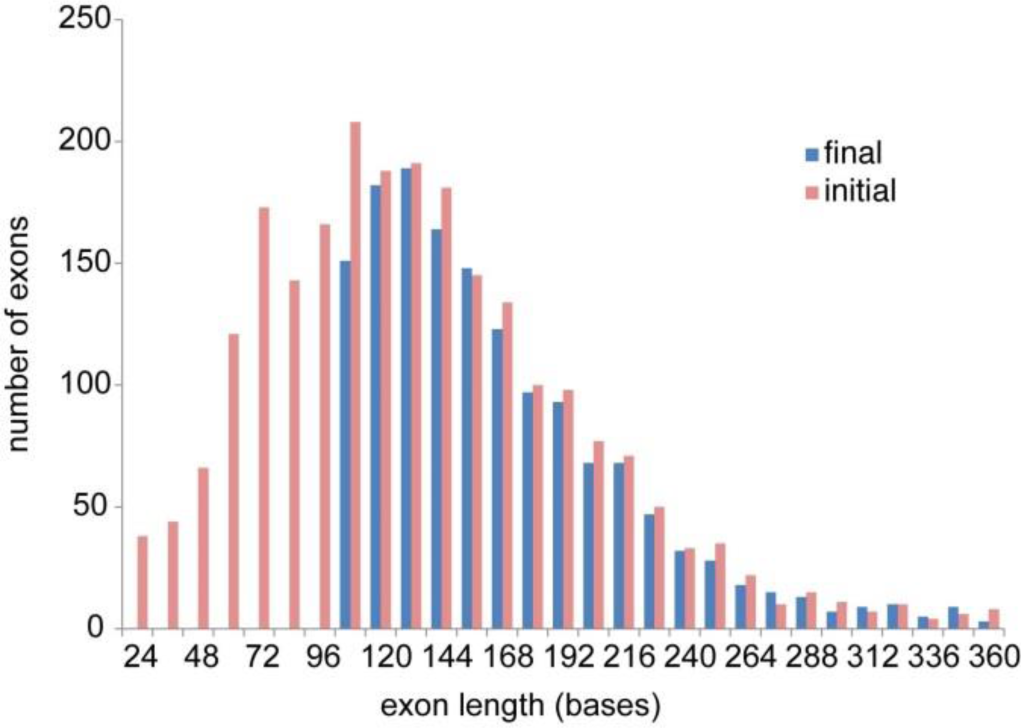
Exon size distribution before and after selection of target.

Across the Ophiuroidea most exons exceeded the hybridisation efficacy limit of 12% genetic distance between probe and target (Hancock-Hanser *et al.* 2013) (fig. 2A). Therefore we included multiple versions of each probe to span the known diversity. We adopted a phylogenetic approach, by designing sets of probes for each major clade identified from our transcriptome tree (fig. 3). We further reduced potential genetic distance by designing artificial exons to represent a clade (based on the ancestral state, see materials and methods) rather than selecting one of the constituent species as an exemplar. We empirically determined that 20 representatives were needed to keep the majority (>80%) of probe distances to transcriptome exemplars to within 12% (fig. 2B & fig. 3). These 20 super-references (SR) were then combined into four 20,000 MYbaits (http://www.mycroarray.com) probe kits of five SRs each, based on the transcriptome phylogeny (fig. 3), in order to be able to flexibly match target samples to their phylogenetically nearest probe sets.

**Fig. 2.**
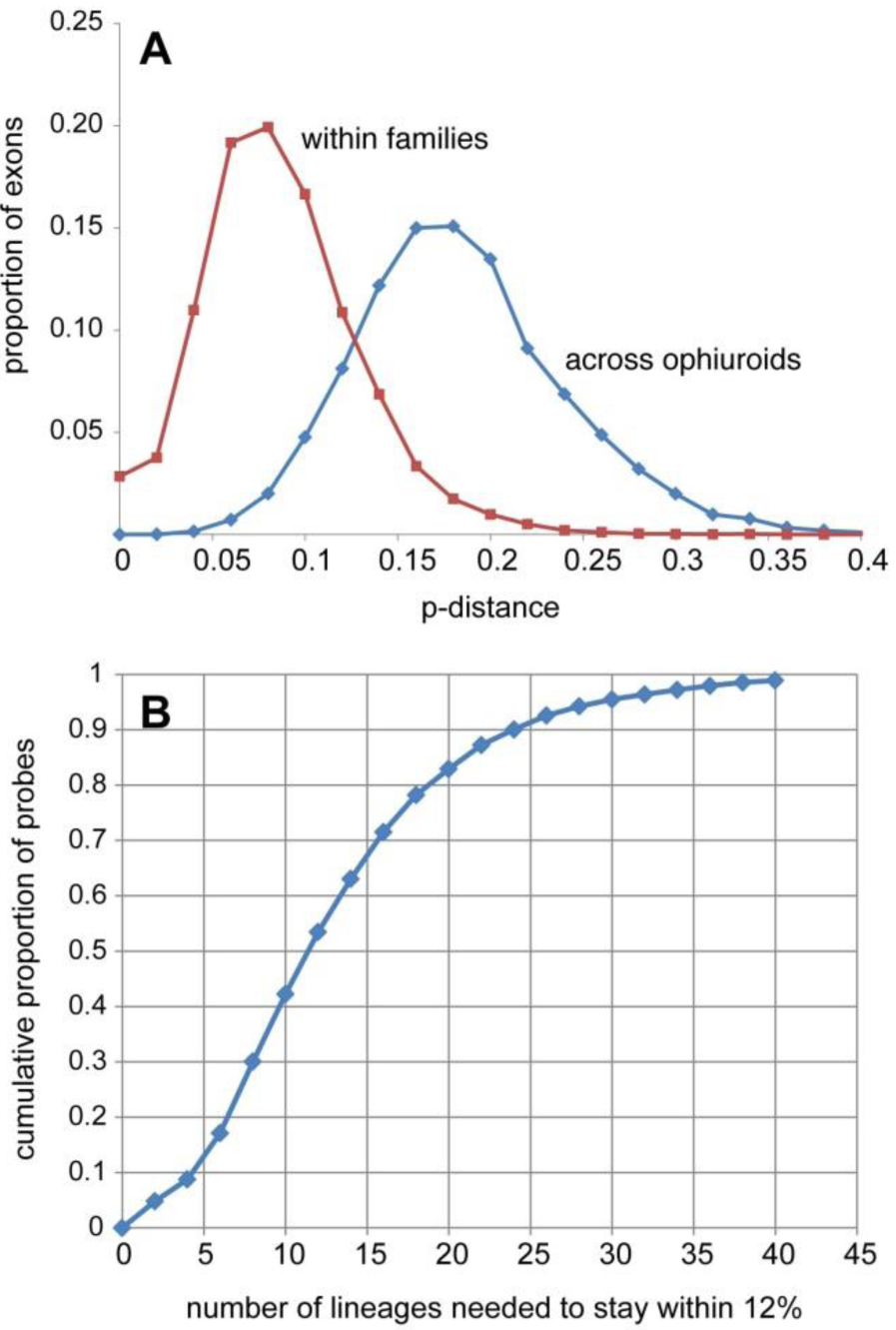
The scale of the problem. **A:** Exon distances among ophiuroids. Across the class most distances are well over the 12% benchmark, within families most are within 12%. **B:** Diversity of capture probes required. The plot shows the cumulative distribution of the proportion of hybridization probes requiring a given number of representatives to ensure that no transcriptome sequence is more than 12% different. With 20 lineages 83% of probes fall within this limit across the candidate target of 425 genes.

**Fig. 3.**
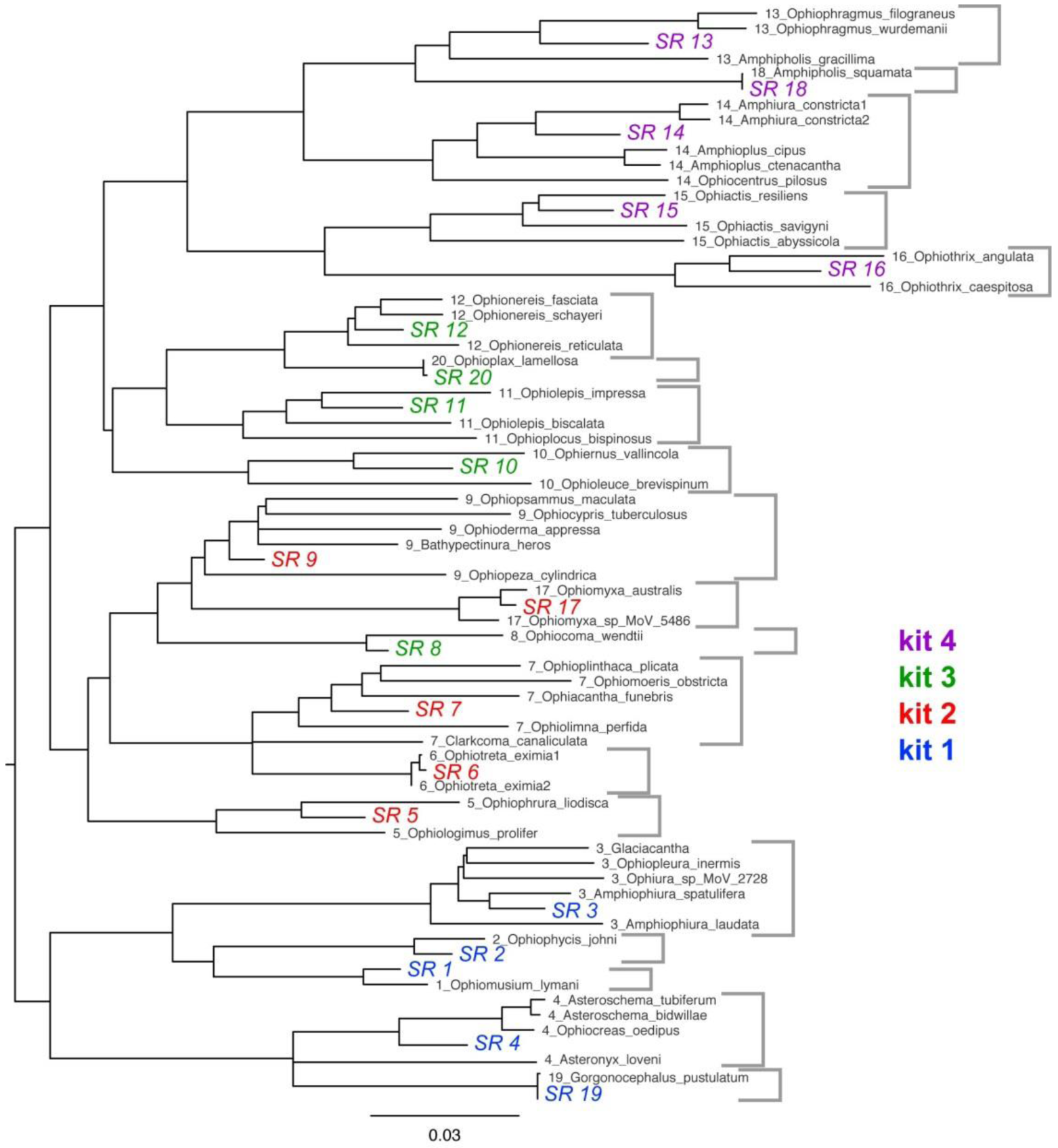
Simple p-distance neighbour-joining tree of the aligned 1552 exons of the original 52 transcriptome taxa and the subsequent 20 representative sequences (labelled “SR”). These are colour-coded by the four kits into which they were combined. Taxa contributing to a representative are indicated by the first number in the label and grey brackets to the right. Note that kit 3 contains the clade 8 representative even though it is phylogenetically closer to the kit 2 sequences. This tree is essentially the same as the full transcriptome analysis in O’Hara et al. (2014b).

### Sequence recovery

To reconstruct exons we mapped reads using two custom pipelines in order explore the issues in creating phylogenomic datasets across an entire class of marine invertebrates: 1) Direct mapping against the best matching member of the 20 representative SR sequences used as the basis of the hybridization probes, and 2) mapping reads to a sample-specific super-reference (ASR) derived from TRINITY *de novo* assembled contigs mapped against the SR set using amino-acid translations (see methods and materials for details).

We obtained usable exon-capture data from 365 samples (Table 1, Supplementary Table S1), with median 0.89 million individual trimmed reads (fig. 4A). Direct SR mapping returned median 45% reads on target (fig. 4B) and coverage per million reads of 172 (fig. 4C). More importantly, the variance in per exon and per sample coverage was reasonably low (SD/mean <1; figs 4C, 5) such that the proportion of sites of the whole target (285,165 sites) with coverage greater than our cut-off limit (>4) averaged 0.93 (fig. 4D). Across all our samples, each SR was selected at least once, with sample target to closest SR p-distances averaging 4.5%, up to a maximum of 11.4% (fig. 4E). The sample-specific ASR mapping returned very similar overall results [Table 1; for clarity not plotted on fig. 4] but with slightly less heterozygosity (0.0065 versus 0.0073) and slightly higher distance to the closest SR (0.050 vs 0.045). Across the core test sample set, 95% of exons had at least ¾ of sites with coverage >4 but 37 exons (2% of the target sites) were never recovered irrespective of distance to the SR or number of reads on target. The common factor here appears not to be variability but that almost all had sections of at least one of the 20 references masked by the MYbaits screening algorithm, indicating that they were problematic to begin with, typically containing quasi-repetitive motifs.

**Fig. 4.**
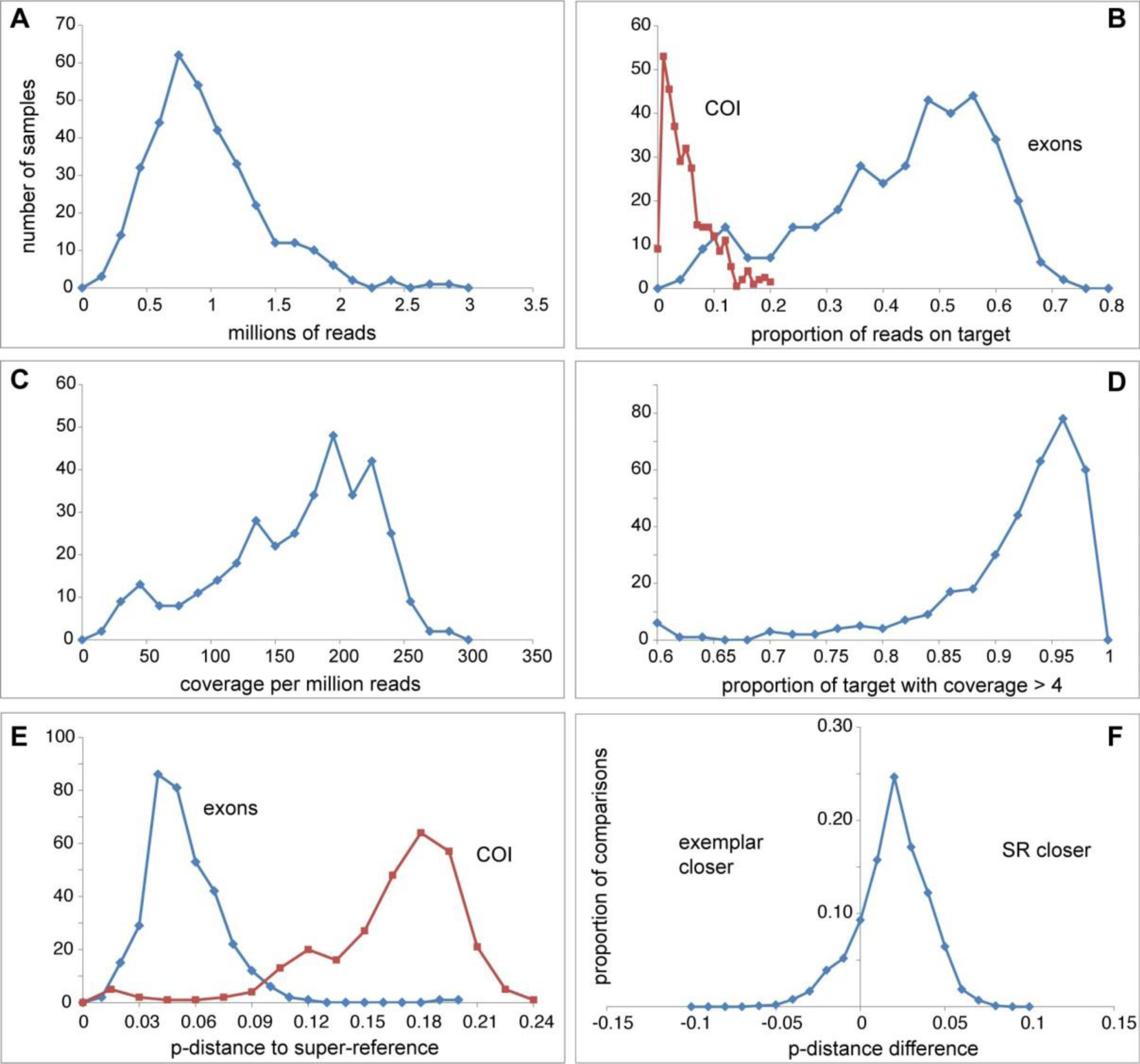
Summary of exon-capture and direct mapping performance. The plots show the number of samples (y-axes) for five key statistics A-E (x-axes). Blue and red refer to exons and COI (B & E). The sixth plot (F) shows proportion of comparisons (y-axis) against the difference in p-distance between sample and SR and sample and transcriptome exemplar, for clades with more than one exemplar (see figure 3). Assembly-based mapping results appear much the same.

**Fig. 5.**
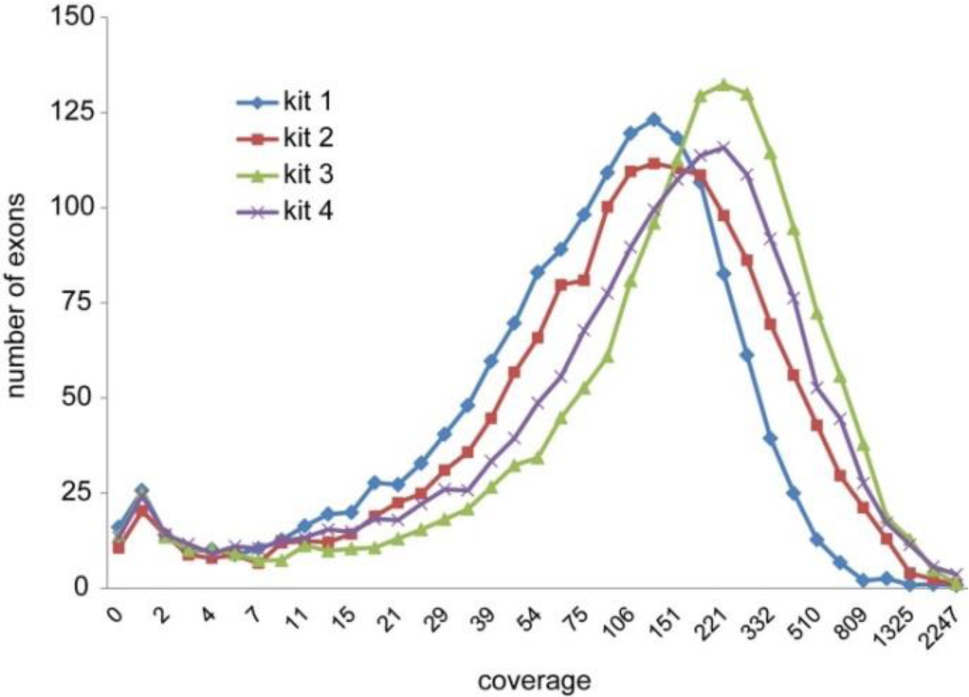
Exon coverage distributions across a 44-sample test set. The four kits are colour-coded as per figure 3.

**Table 1.** Summary of exon-capture performance by kit.

| kit | n | reads | p-RoT | coverage | cov/Mr | p-cov | p-cov>4 | het |
| --- | --- | --- | --- | --- | --- | --- | --- | --- |
| 1 | 109 | 0.74 | 0.41 | 117 | 155 | 0.900 | 0.865 | 0.0071 |
| 2 | 100 | 0.86 | 0.42 | 139 | 160 | 0.949 | 0.911 | 0.0065 |
| 3 | 95 | 0.99 | 0.39 | 153 | 150 | 0.917 | 0.870 | 0.0100 |
| 4 | 61 | 0.99 | 0.46 | 174 | 171 | 0.927 | 0.891 | 0.0118 |
| wildlife | 32 | 1.09 | 0.36 | 159 | 136 | 0.930 | 0.893 | 0.0084 |
| ASR | 365 | 0.89 | 0.42 | 145 | 159 | 0.928 | 0.899 | 0.0065 |

Four single kits and “wildlife” all-four-kits-in-one using direct SR mapping: number of samples; reads in millions; proportion of reads on target; average coverage; average coverage per million reads; proportion of target hit; proportion of target with coverage >4; proportion of sites that were polymorphic; ASR assembly-based mapping results.

To assess the value of using artificial SR for mapping, we compared genetic distances between sample and closest SR, and sample and individual transcriptomes, for clades where we had created a SR from more than one transcriptome species. Overall, for 79% of comparisons, the artificial super-references were closer than the transcriptomes (fig. 4F), and in all but clade 4 (which were drawn towards a rarely sampled divergent lineage) the majority of SR distances were closer than any transcriptome. The artificial SR have additional advantages in trading increased distances to close samples for reduced distances to divergent samples, and because all of the transcriptome exemplars had missing data that must be filled anyway. Thus creating artificial exon sequences directly helped mapping, and by inference may have helped capture.

There was a strong relationship between distance to closest SR and proportion of target recovered that fits a quadratic polynominal function (fig. 6; R^2^ >0.8), such that beyond a certain distance the proportion of target recovered with adequate coverage rapidly falls away. There are two parts to this: 1) failure of the probes to capture the sequence in the first place; 2) the target is too distant from the reference to directly map. In this extensive set of samples only a handful appeared to fall near or beyond these critical limits, a matter we investigated using the assembly-based mapping, which should more represent probe capture limitation alone. This returned a shallower target recovery function (fig. 6 blue line) with substantial gains of 10-20% for the most divergent samples, especially within the more variable exons (gain in coverage versus exon variability correlation coefficient=0.29), resulting in a yet greater gain in information and hence the slightly higher distances to the closest SR. Nevertheless, target recovery for these samples was still well below the overall average of ~90%, highlighting the limits of the single-kit probe capture.

**Fig. 6.**
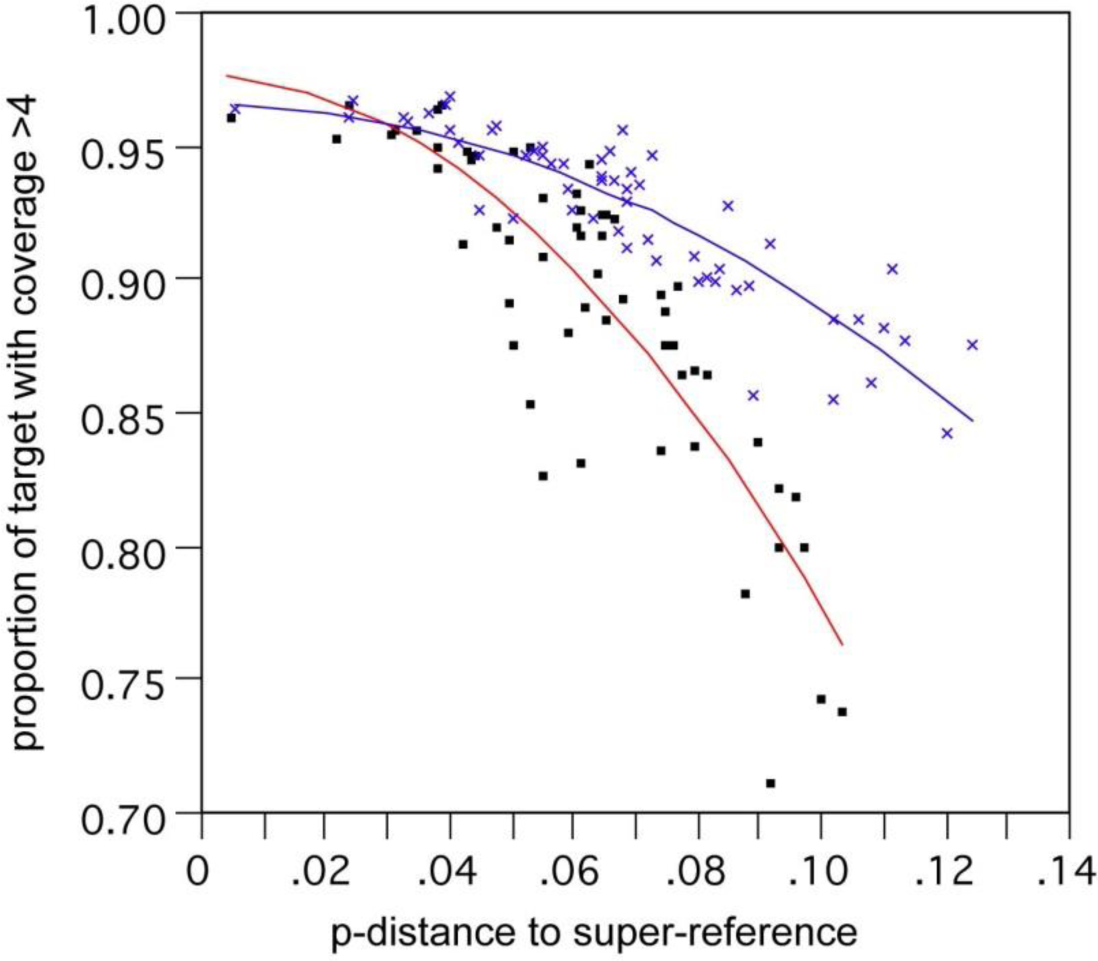
Correlation of proportion of target recovered versus distance from super-reference. Analysis based on a test subset of 59 samples. Lines show two-degree polynomial best fit for direct mapping (red line, black squares) and sample-specific assembled SR mapping (blue line, blue crosses).

### ‘Wildlife’ capture

The bulk of the samples were successfully recovered using one of the four probe kits, however not all taxa can be assigned *a priori* to a kit, or may be divergent from all kits. Therefore we tested exon-capture using all four kits combined (dubbed ‘wildlife kit’ captures), on a subset of samples previously hybridised with one of the four 5-lineage kits. Results were essentially as good as the single-kit captures (Table 1), with ratio of wildlife/single kit coverage averaging 1.00 and none less than 0.86, indicating that high probe diversity and concentration is no impediment. More importantly, one poorly recovered highly divergent sample *(Ophiomyces delata* BP34) now returned 13 times greater proportion of reads on target and three-times the target recovered (from 0.22 to 0.65).

### Polymorphism, paralogy and exon boundaries

Overall there was on average 0.7% of sites with two base states (Table 1), and negligible sites with more than two states (average 6, maximum 73 per sample, out of 285,165 sites). The distribution of the putatively heterozygote sites among exons was largely in keeping with coalescent exponential expectations of allelic divergence (fig. 7). However, 20 exons consistently showed an excess of polymorphic sites across the test sample set, indicative of being confounded by closely related loci, for example pseudo-genes. Removal of these (amounting to 1.5% of the target) largely corrected the observed distribution.

**Fig. 7.**
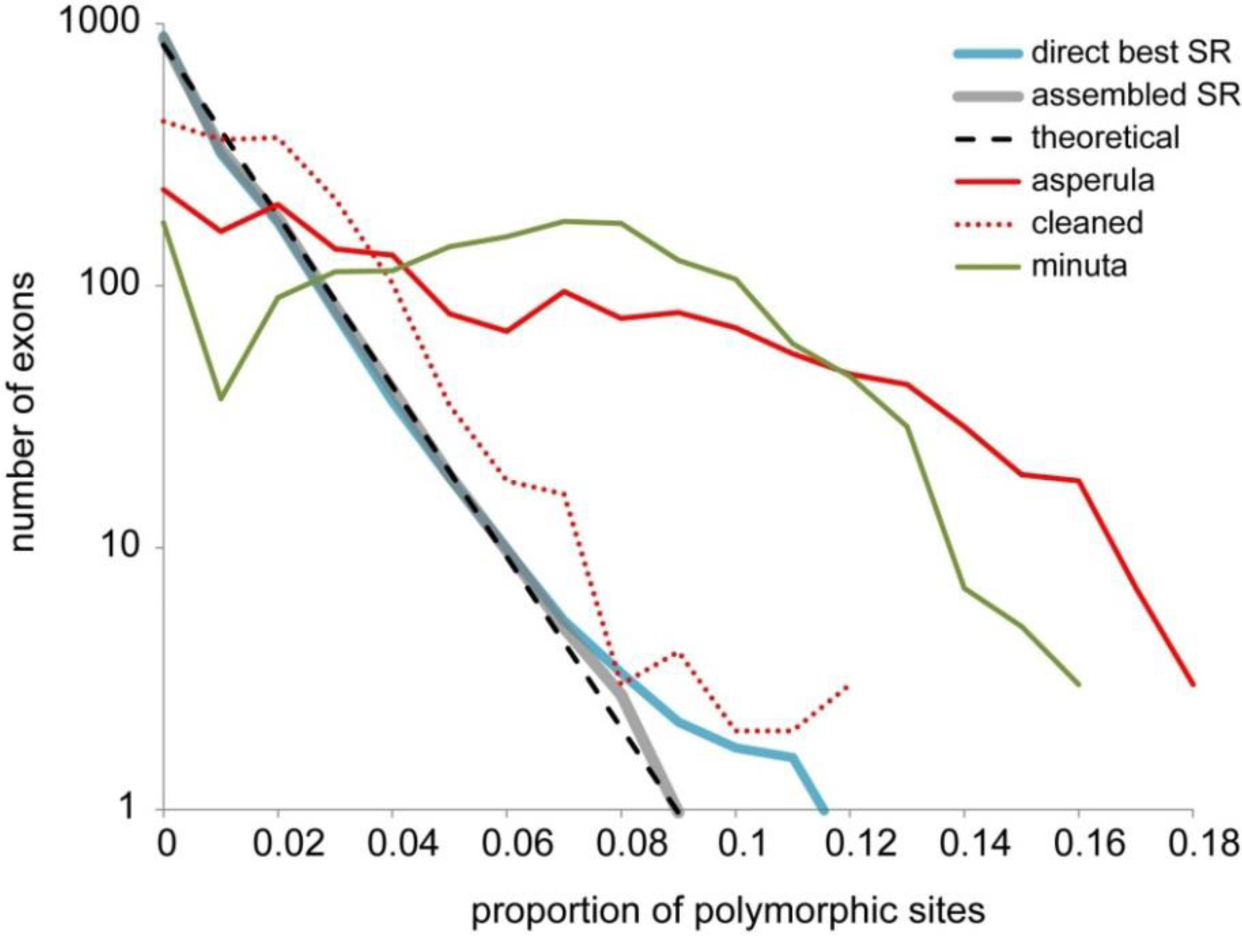
Distribution of exon polymorphism. The thick lines show average distribution of proportion of polymorphic sites per exon across a 44-sample test set: blue direct mapping, grey assembly-based mapping. The dashed line shows the log-linear coalescent expectation. The red lines show *Ophiactis asperula* F167536 before and after filtering of contaminating reads. The putative polyploid/hybrid *Amphistigma minuta* F173962 is shown in green.

In addition to unexpected paralogs, two other problems caused elevated polymorphism: cross-contamination and genuinely divergent alleles (fig. 7). One example is *Ophiactis asperula* that was (inadvertently!) badly contaminated with a divergent species (identified by TRINITY assembled COI contigs) resulting in 6% polymorphic sites, spread across most exons. Filtering the reads of the offending contaminant cleaned this particular sample enough to be used in phylogenetic inference (fig. 7) but we were forced to discard several other contaminated samples that could not be adequately filtered. The second example, *Amphistigma minuta*, is a possible polyploidy or hybrid, where only one COI contig was recovered but at least 90 percent of exons had ~7% polymorphic sites. Phylogenetic assessment of TRINITY contigs of six example exons showed in each case two or three distinct variants belonging to the same lineage in the Amphiuridae (SR 18; fig. 3), accounting for the mapped polymorphic sites.

The exon boundaries in our SR set were based on the half-billion year distant *Strongylocentrotus* genome but ~80% conservation with the sister phylum *Saccoglossus* suggested that ophiuroids would be quite similar. Nevertheless, analysis of read match ends across 44 test samples indicated there were at least 63 substantial differences to our *a priori* boundaries shared across all major ophiuroid lineages. In at least 20 instances adjacent exons were actually contiguous, that is, introns were absent. Given the large phylogenetic scale there could also be some boundary differences among ophiuroid lineages. This remains to be fully assessed but examination of the test samples showed 21 consistent differences between clade 12 and clade 15 taxa. Broadly, using the criterion of a fixed difference in all samples in any one of five major lineages across 44 test samples, 7.4% of exons contained some boundary change.

### Mitochondrial COI gene

Due to much higher divergence levels, the COI exon-capture sequences were determined from TRINITY assembled contigs. Much higher divergences potentially posed a problem for capturing COI but this appears to have been ameliorated by the substantially greater natural abundance of mitochondrial DNA (estimated from unpublished ophiuroid partial-genomic data at ≥100-times that of exons). Empirically, recovery of the COI gene was quite variable (fig. 4B) and in some cases (n=9) failed entirely but on average accounted for 5% of all reads, resulting in a high median coverage of 3000. Distances to the closest reference for COI were frequently well beyond the 12% level (fig. 4E) however there was useful enrichment of on-target COI (red line) versus off-target flanking mtDNA sequences (fig. 8). Similar analysis of six samples, indicated that COI enrichment was an order of magnitude less than nuclear exons, which we estimate to be several thousand-fold.

**Fig. 8.**
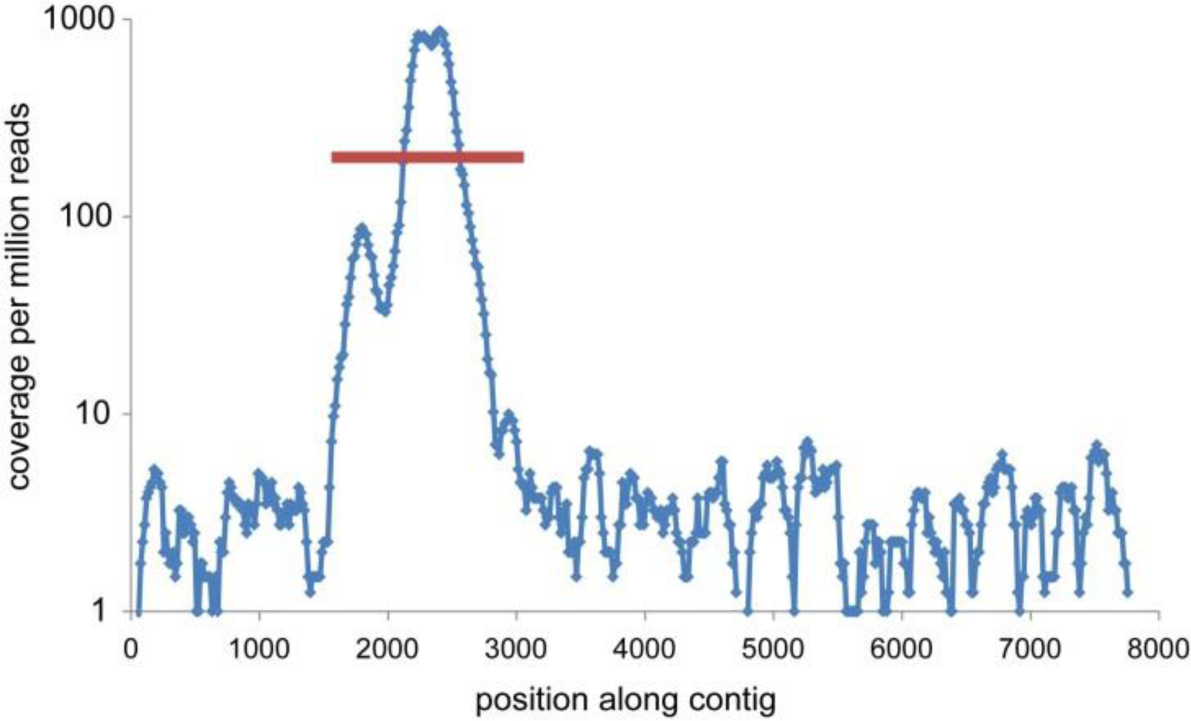
Mitochondrial COI gene capture. Coverage along a long Trinity-assembled mtDNA contig containing the targeted COI gene (red line).This test sample *(Ophiotreta valenciennesi* UF8999) had 4.4 million reads and the COI gene was 17% different to the closest reference.

### Limits of precision

An important criterion in exon-capture is accuracy: how close is the mapped sequence to the true genotype? We do not have a known genotype but gauged consistency in two ways: by comparing replicated captures, and by comparing pipelines for the same capture. The first approach used nine phylogenetically diverse samples captured twice: once with a single kit and again with the wildlife kit. The second used a different set of 12 samples. For direct SR mapping, replicate captures gave near identical results (average 7 fixed differences) reflecting the deterministic mapping process (Kent 2002). For sample-specific ASR mapping, replicate captures showed higher differences (average 131 fixed differences, or 0.00046) reflecting the more complex assembly-based process (Grabherr et al. 2011). Differences among mapping pipelines (direct, ASR and BFAST/VARSCAN) for the same capture, similarly were around 0.0005 fixed differences. Considering all states, fixed and polymorphic, all types of comparisons gave higher differences of around 0.002. None the less, such values amount to Phred scores of >27, higher than typical read sequence quality cut-offs and as good as most Sanger sequencing. About 11% of differences were associated with exon boundary and indel sites, which only make up 3.5% of all sites, suggesting they have higher error (Homer et al. 2009; Bi et al. 2012).

### Phylogenetic analysis

*Post-hoc* analyses showed that about 3% of our target was not captured and a further ~2% confounded by paralogy, leaving 95% as reasonable data for phylogenomic analysis. Exclusion of these sites gave a 91% (range 41–98%) character-complete data matrix of 275,352 sites by 417 tips, including the original 52 transcriptomes, and covering 380 species in 121 genera of ophiuroids. To assess computational load, a smaller dataset of 239 tips was analysed first: a complete RAxML –f a command of GTR-CAT model fast bootstraps followed by a full ML search was run (via CIPRES) taking 2660 CPU hours. This result was then compared to a tree made from the all-compatible consensus topology of an independent set of 100 GTR-CAT fast bootstraps, and ML branch length optimization only of this consensus topology. This process took 220 CPU hours and returned an identical tree – same topology, support and branch-lengths as the full search but taking one tenth of the CPU time. Hence this approach was applied to the full 120 megabyte 417 tip data matrix, using 200 bootstraps and taking 550 CPU hours. The resulting tree shown in fig. 9 is highly resolved with 90% of nodes having 100% bootstrap support, including nearly all major lineages. In both the 417 and the 239 tip trees, the sub-tree of the 52 transcriptome species is topologically identical to the original transcriptome tree in O’Hara et al. (2014b). To investigate the consistency of the data, we compared RAxML analyses excluding exon boundary and indel site codons (11% of sites), and using the sample-specific ASR mapping data. Excluding exon boundary and indel codons made negligible difference (topology differed for only three minor nodes with <60 BS support). The two mapping pipeline dataset trees had 15 differences, mostly non-significant intra-specific tips. The exception was the position of the divergent genus *Ophiopsila* which did differ by one major node (with 100% BS) in the direct mapping data tree (fig. 9 Ophionereididae, SR20 group, marked with a cross).

**Fig. 9.**
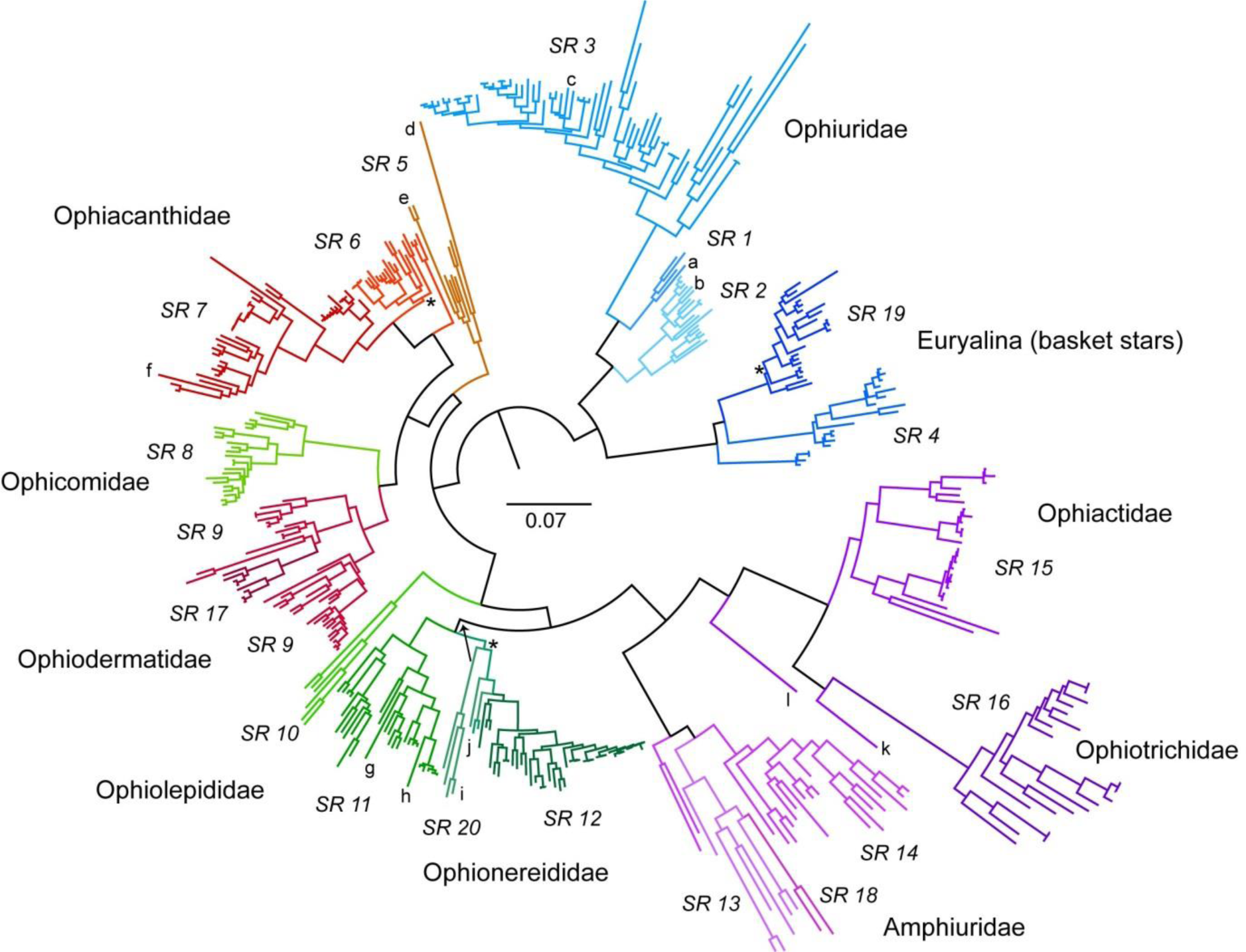
An ophiuroid exon-capture phylogeny. The tree is a RAxML codon-position GTR-CAT fast bootstrap consensus with ML branch lengths, for 417 samples (380 species) from 275,352 sites in 1490 exons, rooted according to O’Hara et al. (2014b). Centre bar indicates divergence scale; taxon names have been omitted for simplicity; lineages are colour-coded and labelled by mapping SR, and some large family groups indicated. The three higher-level nodes with bootstrap support <95% are marked by asterisk; major difference between mapping pipelines in placement of the SR20 lineage indicated by arrow. Genera mentioned in the text are denoted as follows: a - *Ophiomusium*, b - *Astrophiura*, c - *Ophiosparte*, d - *Ophiomyces*, e - *Ophioscolex*, f - *Ophiocanops*, g - *Hemieuryale*, h - *Astrogymnotes*, i - *Ophiopsila*, j - *Amphilimna*, k - *Ophiopholis*, l - *Ophiothamnus.*

## DISCUSSION

We have presented a summary of our universal class-wide exon-capture system, designed principally to streamline tree of life-scale phylogenomics. Our exon-capture system reliably sequenced ~90% of 1552 exons from hundreds of species from across an entire class of marine invertebrates, the Ophiuroidea. Only a handful of samples appeared to have been too divergent to capture and map fully (e.g. the distribution tails in figs 4D, E; fig. 6).

### Multiple lineage capture system

Compared to most hybridization enrichment phylogenetic studies published to date, our system has both a large phylogenetic scale and a high target recovery (c.f. Bi et al. 2012; Faircloth et al. 2012; Lemmon et al. 2012; Hedtke et al. 2013; Leaché et al. 2014; Mandel et al. 2014; Tilston-Smith et al. 2014)). The major aspects contributing to this consistent recovery would be 1) use of multiple references based on a thorough transcriptome phylogeny of the class (O’Hara et al. 2014b) 2) excluding exons and parts of exons that would be difficult to capture or map or align; 3) designing our probes and references from synthetic ancestral-state derived sequences; and 4) mapping pipelines designed to maximize the recovery of captured exons. Use of long 120-base RNA in-solution hybridization probes may have also helped. Final target size of 285kb in 1552 exons is relatively modest, limited to transcripts commonly expressed across a class and filtered to a tractable target. However the benefit of this *a priori* trimming of the target, in combination with multiple phylogenetically diverse references, means that the final output is dense in information with a high proportion of useable exons and variable sites.

Having an *a priori* good measure of the phylogenetic diversity of the target allowed us to create multiple artificial consensus representative sequences for probe design and for mapping references. Our procedure for creating these was somewhat *ad-hoc* and doubtless could be refined but the key point is that the concept appears valid, narrowing the range of distances to novel samples to improve capture and mapping. We combined a 12% rule of thumb, measures of diversity across the class and MYbaits technical requirements, to estimate that 20 representatives would be sufficient to capture class-wide phylogenetic divergence. Our target recovery (figs 4, 5) indicated that this level of replication was more than adequate for probe capture (with the exception of some mitochondrial COI) and direct mapping of sequenced exon reads. While we cannot calculate exactly how much the artificial SR actually helped in target recovery, applying the ASR coverage versus distance function (fig. 6 blue line) to the sample to the distances between sample and reference underlying fig. 4F would imply up to 40% more missing data using single-species exemplars instead of artificial SR.

### “Wildlife kit”

There appears to be little biochemical impediment to combining multiple homologous probes into the one bait kit. For the majority of samples, we used probe kits containing five related lineages. However, we also combined probes from all 20 lineages to create a ‘wildlife kit’ potentially capable of hybridising to any ophiuroid species. While we only tested this on a small number of samples, the results were encouraging. There was no degradation in the proportion of exons recovered using the wildlife compared with the 5 lineage kits on the same libraries from 24 test samples. Moreover, for one divergent species, *Ophiomyces deleta*, the proportion of exons reliably returned rose from <30% to >60% using the wildlife kit, and also improved somewhat for other taxa.

### Multiple lineage mapping system

The next key issue is how to faithfully reconstruct clean homologous informative sequences from divergent exon-capture data. While multi-transcriptome phylogenetic analysis aided in the *a priori* exclusion of problematic loci (e.g paralogs) that are too close to be reliably distinguished from the desired target, not all such loci are necessarily accounted for, such as infrequently expressed genes and in particular pseudogenes. Neither do transcriptomes provide information on exon boundaries. While a genome sequence does account for all loci and exon boundaries it only does so for that genome, and across any substantial phylogenetic diversity there are likely to be lineage-specific paralogs (e.g. pseduogenes) and exon boundary shifts (Lynch 2002; Zhang 2003; Parmley et al. 2007; Roy 2009). Therefore some *post-hoc* target filtering is necessary. The problem with lineage-specific paralogs is that unless they lie outside the reference clade they cannot readily be distinguished from the true ortholog by match similarity to the reference: they will in effect appear as equidistant competing versions. Sample cross-contamination, a constant danger in any laboratory work, can pose a similar difficulty. Therefore it is important to have a mapping system that identifies problematic exons and samples (Lemmon and Lemmon 2013; Mandel et al. 2014).

Hence we developed two read mapping systems that have a number of features worth mentioning. They are pitched at tree of life-scale phylogenetic analyses and so what little alignment is required was built into the mapping, allowing new samples immediately to be slotted into existing datasets. Because reads are clipped to match length, mapping was resilient to unknown target exon boundaries. The direct SR mapping was fast and consistent, at the expense of loss of data in divergent samples, and higher interference from close paralogs and (relatively abundant) contaminants resulting in ambiguous sites. The ASR mapping is more complex and somewhat less consistent but recovers more target from divergent samples. It has less interference from close paralogs and (relatively abundant) contaminants due to the tighter re-mapping criteria (7% versus 14%).

The key point is that, in both methods, the relaxed mapping makes a virtue of necessity by flagging problematic exons and samples via the pattern of elevated polymorphic sites, which can then be used to diagnose potentially confounded exons and contaminated samples (fig. 7). This also has advantage in highlighting taxa that may truly have complex divergent allele patterns (e.g. polyploids). For phylogenetic analysis, after excluding certain exons (and samples!) entirely, coding remaining polymorphic sites as ambiguous is a reasonably conservative approach. Both of our mapping systems probably have some inconsistency in mapping around indels (Homer et al. 2009). However as we eliminated most of these in the target *a priori*, the remainder comprise a tiny fraction of sites and are better off excluded in subsequent analyses than to try improving their alignment. Similarly for sites immediately adjacent to exon boundaries.

Altogether, these pipeline attributes are desirable for tree-of-life scale phylogenomic datasets, where trying to account for idiosyncratic indels, exon boundaries and confounding paralogs in thousands of loci for hundreds or thousands of taxa across multiple divergent lineages would be costly for little gain. Being able to add new taxa, and mix and match taxa and exons, without any post-mapping alignment and site filtering, is a great advantage as we continue to gather samples incrementally.

### Refinement and precision

With information on many lineages, we are now in a position to refine the target, revise exon boundaries, and possibly expand the set of super-references via phylogenetic ancestral state inference. For the phylogeny presented here we have only excluded ophiuroid-wide null and paralogous exons but this process could be extended further to a more detailed taxon-by-exon filtering of unreliable exons (e.g. Mandel et al. 2014). For example, across the test sample set the ASR typically had 0.008 sites with competing contigs, falling to 0.005 polymorphic sites in the remapped sequence. Elimination of exons with >0.04 multi-state sites in both ASR and re-mapped sequence excludes 0.003 sites, essentially the previously identified 20 paralogous exons.

Our system is primarily focussed on large tree of life-scale phylogenetics, especially of groups with little genomic and phylogenetic information. However, the exon set appears to contain useful information at much finer phylogenetic scales. For example, across our samples, 25 independent close intra-specific pairs typically had 500 fixed-differences in 250 exons. While our estimate of consistency would suggest an ‘error rate’ of around 150 fixed-differences, this potentially still leaves several hundred SNPs for multi-locus species-tree and historical demographic inference (Knowles 2009). For these purposes data would be re-mapped with appropriately tighter stringencies and coverage limits, along with separating alleles and including genotype quality likelihood metrics (e.g. Altmann et al. 2012).

### Mitochondrial COI

We included the mitochondrial COI gene in our probe design to help verify the sample identity against ‘barcode’ datasets, identify contaminants and facilitate the future incorporation of legacy COI sequences from other species into our phylogeny. Although the ~50% off-target reads will contain mtDNA, the probes were essential in recovery of the COI (fig. 8), typically with coverage far in excess of what is needed but with little apparent decrement to exon capture (correlation coefficient between relative abundance of COI reads and nuclear exon coverage =−0.043, p=0.43). The major concern for COI is the shear diversity of sequences that are recovered, indicating all manner of minor cross-contamination as well as the expected miss-indexing. This requires a significant amount of effort to interpret. As the best matching or highest coverage contigs were not always the correct one, all candidate contigs needed to be included in phylogenetic assessment of orthology. Furthermore, the TRINITY software occasionally did generate chimeric sequences from mixtures of divergent samples (Grabherr et al. 2011). Despite these complications, our experience has been that having a tool to validate sample identity is highly desirable.

### Ophiuroid Phylogeny

Generation of phylogenetic trees from massive genetic datasets is problematic. While recent methods are remarkably efficient (Aberer et al. 2014; Stamatakis 2014), genomic-scale tree of life phylogenies still require vast amounts of CPU time. On the other hand, such large datasets should be very powerful and contain a great deal of information on sequence evolution patterns. Therefore they ought to be well-suited to the very efficient GTR-CAT approximation, allowing a great reduction in CPU time for very little loss of inference (Stamatakis 2014). For our dataset, the fast approximation returned the same tree as the full search. Moreover a test set of just 34 exons (using full RAxML analysis) returned the same tree topology, with 76% of nodes with 100% bootstrap support, from one-tenth of the sites (data not shown). Conversely even colossal amounts of data may not resolve all phylogenetic questions (e.g. Jarvis et al. 2014). Most of our tree discrepancies involved tip intra-specific complexes, which are better analysed in a multi-locus coalescent framework (Knowles 2009). But for very large phylogenetic analyses a concatenated approach is sensible. The tree presented in fig. 9 contains 417 tips covering 380 species in 121 genera across all currently named ophiuroid families with full support for 90% nodes, and would be one of the most powerful of any metazoan class published to date. It is fully consistent with the transcriptome tree but not congruent with the current classification of the Ophiuroidea (O’Hara et al. 2014b) nor with any previous published hypotheses of super-family groups. Hence the results make possible a whole-scale taxonomic revision of the entire class, involving detailed interrogation of phylogenetic hypotheses and mapping of morphological characters. Such a task is not possible or appropriate here but below we draw attention to several major aspects.

### Taxonomic implications

At the highest level our phylogeny forms two clades (fig. 9). The first includes the Ophiuridae, the generic-complex *Ophiomusium/Ophiosphalma/Ophiolipus* (formerly in the Ophiolepididae), and the basket stars (euryalins). The second clade includes all remaining families. This separation was first suggested by Martynov (2010) who noted that the euryalids and ophiurids have similar primitive arm spine articulation morphologies and that all other families “form a compact group with numerous intermediate taxa”. However, far from being a homogenous group, we find that this latter assemblage comprises two ancient subclades, one including the Ophiacanthidae, Ophiocomidae, Ophiodermatidae and ‘ophiomyxids’, the other containing the Ophioleucidae, Ophionereididae, Ophiolepididae, Ophiactidae, Ophiotrichidae and Amphiuridae. These grouping are not consistent with any previous classifications (O’Hara et al. 2014b).

Many existing families and genera are polyphyletic (Ophiolepididae, Ophiomyxidae, Ophiocomidae) or paraphyletic (Ophiacanthidae, Ophiodermatidae, Ophiactidae). Thus many characters that have been used to define higher-level taxa are evidently homoplastic, including the reduction of the external skeleton, the form of the arm vertebrae, and the position of oral papillae and tentacle pores on the jaws. Only microscopic characters of the lateral arm plates, most notably the form of the articulation with the arm spines, appear to be reliably diagnostic for family-level clades (Martynov 2010; Thuy and Stöhr 2011; O’Hara et al. 2014b).

Our phylogeny also resolves the position of many controversial taxa. *Hemieuryale*, type genus of the Hemieuryalidae, falls within the Ophiolepididae, a relationship completely at odds with its traditional taxonomic placement which emphasised the form of the arm vertebrae. Surprisingly, *Astrogymnotes* formerly considered an ophiomyxid is also a derived member of the Ophiolepididae. *Ophiocanops* is an aberrant ophiacanthid, related to *Ophiomoeris.* Due to some unusual features such as the presence of gonads under the dorsal arm surface of some species and connective tissue covering the ambulacral groove, the genus has been previously classified in its own family (Mortensen 1932), as a relict stem ophiuroid (Fell 1963), or as an ophiomyxid (Stöhr et al. 2008). The pentagonal *Astrophiura*, originally considered close to the asteroids (Sladen 1889), forms a distinct clade with similar genera *Ophiomisidium* and *Ophiophycis*, that is sister to the Ophiuridae *sensu stricto.* On the other hand, the large Antarctic carnivore *Ophiosparte gigas*, previously classified as an ophiacanthid or ophiomyxid, falls well within the Ophiuridae, in a soft-skinned group including *Gymnophiura, Ophioperla*, and *Ophiopleura. Ophiomyces, Ophioscolex, Ophiopsila, Amphilimna, Ophiopholis* and *Ophiothamnus* each form the basis of divergent lineages that appear to be deserving of family-level status. A detailed systematic analysis is in preparation.

The data generated through emerging next-generation technologies will not only resolve contentious phylogenetic problems but also provide a solid basis for evolutionary, biogeographic and conservation studies. Marine invertebrate taxonomies to date have been too uncertain or unresolved to be useful in such analyses. Many genera such as the gastropod *Conus*, the galatheid *Munida* and the ophiuroid *Amphiura* have hundreds of species (WoRMS Editorial Board 2015) spread across the globe, providing poor phylogenetic resolution. O’Hara et al. (2014b) and the current study provides clear evidence that historical qualitative taxonomic diagnoses can be a poor guide to phylogenetic relationships. Large phylogenomic datasets combined with recently assembled global distributional data (OBIS 2014) will be a powerful tool to explore the origin and distribution of marine life.

## MATERIALS AND METHODS

### Target selection

We identified exons from a 425 nuclear gene alignment (O’Hara et al. 2014b) assembled from 52 ophiuroid transcriptomes, six outgroup transcriptomes and three outgroup genomes (the fish *Danio rerio*, hemichordate *Saccoglossus kowalevskii* and echinoid *Stronglyocentrotus purpuratus).* We also included the mitochondrial COI gene (also derived from the transcriptomes). We selected these genes after reciprocal BLAST and phylogenetic assessment of orthology, and presence in at least two-thirds of the ophiuroid transcriptomes (see O’Hara et al. 2014b for details). We estimated exon boundaries by mapping corresponding proteomes against the three included genomes using the program BLAT (Kent 2002). For simplicity all boundaries were made to be in frame. Within these 425 genes, 83 percent of exon boundaries appear to be conserved (within four codons) between *Strongylocentrotus* and *Saccoglossus* and 75 percent between *Strongylocentrotus* and *Danio;* consequently we used the closest genome *(Strongylocentrotus)* as the basis for breaking up the ophiuroid transcriptome data into putative exons. After removing all outgroups, the starting alignment for selecting exon-capture targets comprised 425 genes with 2544 nominal exons spanning 427,832 sites (142,611 codons; fig. 1 “initial”). Of these 2544 exons, we excluded 1036 because they were too short (<99 bp), had excessive length variation or repeat elements, or were missing from several of the major ophiuroid clades of our transcriptome phylogeny (O’Hara et al. 2014b). Some exons were trimmed in length and some (44) were split, to avoid complex regions. This left a final target of 1552 nominal exons in 418 genes spanning 285,165 sites (fig. 1 “final”).

### Accommodating phylogenetic diversity

We investigated two approaches to constructing artificial representatives from selected clades in our transcriptome data. For each base along an exon we 1) selected the most frequently-present nucleotide within the clade, and 2) derived an ancestral state [via accelerated transformation in PAUP 4b10; (Swofford 2003)]. Both approaches can reduce evolutionary-shallow substitutions but the frequency method will push the representative towards the most speciose lineage in the clade, while the ancestral approach will push the representative towards basal lineages. The best option then depends on the sampling of clade diversity. Consequently, we implemented a mixed solution, constructing the final representative exons by randomly selecting bases from both the frequency and ancestral models. Finally, we substituted the phylogenetically closest sequence for any missing data.

We selected the MYbaits (http://www.mycroarray.com) sequence capture system because they offered custom-built kits of long probes (120bp) for relatively small targets (minimum 20,000 probes). We determined how many clade representatives were needed to keep probe distances to all the members of that clade within 12% (fig. 2B & fig. 3). With 20 clade representatives, based on one to five transcriptomes each, 83% of 120 base probes remain within 12%. These 20 representatives were then split into four 20,000 probe kits of five representatives each, based on the transcriptome phylogeny (fig. 3). The major ophiuroid clades derived from our transcriptome data were uneven in terms of their putative species richness and genetic diversity. Consequently one kit contained a probe set from a distant lineage (kit 3, fig. 3). We retained duplicate sequences from different clades (e.g. for conserved or substituted exons) and further duplicated small exons (99-120 bp) to reduce variation in probe concentration across the target sequences. Based on these 1552 target exon sets, final bait sets were designed and synthesized by MYcroarray using 2x tiling.

### Sample selection and Laboratory procedures

DNA was extracted using Qiagen DNeasy Blood & Tissue kits from a diverse range of shallow and deep-water ophiuroid species collected since the year 1999 and fixed/preserved in ethanol (70–95%). We selected several hundred reasonable quality DNA extractions (based on agarose gels), with similar numbers putatively assigned to each of the four probe kits (Supplementary Table). The extractions were dried, stored and shipped to the Georgia Genomics Facility on 96-well DNAStable plates (Biomatrica).

Dried DNA extractions were rehydrated in Tris-EDTA, quantified using a 96-well fluorometer and Sybr Green I (Life Technologies) and, where possible, normalized to 15 ng/μl. DNA was sheared by Covaris S2 then transferred to 96-well plates and processed using the KAPABIOSYSTEMS DNA Library Preparation Kit. After library preparation using a truncated common Illumina Y-adapter stub, a standard dual-indexed sequencing adapter for Illumina sequencing was added using a unique combination of indexed i7 and i5 PCR primers for each library in the 96-well plate. After six cycles of PCR, the amplification was checked by agarose gel electrophoresis and low concentration samples subjected to additional amplification. Libraries were AmpureXP purified and the concentration determined by fluorometry. The final amount of library varied substantially due to sample quality but where possible up to 200ng of each library was combined into pools of 8 individuals for sequence capture.

These library pools were concentrated to a volume of 30μl using Qiaquick PCR Purification columns and then further concentrated to 3.4μl using a centrifugal vacuum concentrator. MyBait probes were diluted with water and used according to the manufacturer’s protocol version 2.2 with the optional high-stringency wash conditions. Briefly, heat denatured concentrated library pools were combined with probes, non-specific and MYcroarray supplied Illumina adapter specific blocking reagents, and hybridized for 40 hours at 65°C. Hybridized probes were captured using Dynabeads MyOne Streptavidin C1 beads, washed 3 times at 65°C with a 1:5 dilution of MYcroarray Wash Buffer 2 and the beads resuspended in 30μl of water. Ten μl of beads were then used in a 50μl PCR reaction with 25μl 2X KAPA HiFi, HotStart ReadyMix and 0.3μM each of the Illumina PCR amplification primers. Amplifications were checked by agarose gel electrophoresis after 12 cycles then cycled for an additional 4 or 5 cycles. After AmpureXP purification, the capture pools were resuspended in 15μl EB (Qiagen), quantified by Qubit fluorometry and size distribution checked with the NGS Fragment Analysis Kit on a Fragment Analyzer (Advanced Analytical).

In general we used a one-quarter dilution of a MYbaits kit per capture. We also trialled mixing all four kits together (creating a “wildlife kit”), using one-eighth dilution. Typically, six captured library pools were combined in equimolar amounts (amounting to ≥48 individual sample libraries) and sequenced on a Miseq Desktop Sequencer using the Illumina Miseq Reagent Kit v2 300 cycles with dual-indexed paired-end 150 cycle settings. To estimate enrichment we compared reads on-target and target length (bp) against an estimate of ophiuroid genome size, calculated from unpublished partial-genomic data of *Ophiactis abyssicola* to be ~2Gb.

### Mapping pipelines

Illumina adapters and low quality read regions were removed using Trimmomatic−0.22 (minimum quality score 25 per 4-base window) (Lohse *et al.* 2012). As duplication levels were ~20%, all reads were used. Exon sequences were then reconstructed by mapping reads to a chosen reference. Our primary pipeline mapped reads directly to the best matching member of the 20 representative “super-reference” sequences used to design the hybridization probes. These super-references (SR) are aligned sets of the 1552 exons incorporating 340 separate indels spanning a total of 1089 sites (333 codons), in order to keep the 20 different representatives in alignment with one another, keeping all exon boundaries and indels in-frame.

All mapping was conducted using BLAT (Kent 2002) built into custom Unix shell scripts to interpret the psl format output. The basic pipeline comprised three parts: 1) identifying the super-reference that best matched the sample; 2) mapping all reads to this super-reference; summarizing the output as a site by character state table (for five states: A,C,G,T, other); and 3) inferring a final consensus sequence (and summary statistics) from this table based on a set of rules. BLAT mapping was done using standard two 11-base tile match initiation and overall minimum identity of 0.86, with minimum match and block size filters. To choose the best matching SR a subset of 50,000 reads were mapped onto a subset of 50 variable exons (3.5% of sites) for all 20 super-references. For base-calling, sites with no coverage were coded as “-”, coverage 1-4 coded as “n”, and above this limit a site state was included if abundance was greater than 20 percent. Two-state sites (nominally heterozygote) were IUPAC-coded, more than two coded as “X”. A minimum coverage of 5 for exons was considered to be sufficient to exclude miss-indexing from affecting base-calling, although a higher limit could be used if heterozygote status was critical (Altmann et al. 2012).

A secondary pipeline involved mapping reads to a sample-specific super-reference built around *de-novo* assembled gene fragments (e.g. Bi et al. 2012; Lemmon et al. 2012; Tilston-Smith et al. 2014). All sample read sets were assembled using TRINITY (Grabherr et al. 2011, default settings). The subsequent set of contigs provides an alternative source of candidate sequences, potentially allowing detection of genes that were captured by the hybridization probes but are too divergent to effectively map directly. Assembly-based references raise the issue of selecting amongst different competing contigs. There are various sources of these: paralogs both divergent and close (outside and within the reference lineage); divergent and close contaminants of both relatively low and high coverage, assembly artefacts and true divergent alleles. Our approach is aimed at avoiding choosing the wrong contig, or discarding exons unnecessarily (c.f. Mandel et al. 2014; Tilston-Smith et al. 2014).

To construct assembly-based super-references (ASR) the TRINITY assembled contigs were mapped (at the translated amino acid level with minimum identity 0.86) to the closest SR to generate a consensus nucleotide sequence, marking sites with more than one state due to competing contigs as polymorphic. A final composite ASR was then derived by replacing missing and polymorphic sites with the corresponding positions in the closest SR, to give a fully defined complete ASR of exactly the same length and alignment as the original super-reference set. Reads were then mapped to this ASR as above but at higher stringency of minimum identity 0.93. This approach allows the gathering of all non-overlapping exon fragments due to gaps in coverage or unexpected exon boundaries. Resolving sites with competing contigs in favour of the closest SR for re-mapping has a number of attributes. It effectively excludes from mapping divergent (out-) paralogs and contaminants, relatively low coverage close contaminants will not contribute to the final consensus, while interference from closely related paralogs and relatively abundant contaminants is retained in the pattern of multistate sites in both the mapped contig ASR and the subsequent re-mapped sequence.

Both pipelines take advantage of BLAT accepting reference gap sites and clipping reads to match length. Thus, because novel insertions are excluded and all the exons in the 20 super-references are pre-aligned, the output sequences are aligned as they are mapped, and can be added to pre-existing datasets without any further processing.

For a small subset of taxa, exons were mapped using BFAST (Homer et al. 2009) to provide BAM format files for visual inspection of exon boundaries in IGV (Robinson et al. 2011). We used the default settings for BFAST with the exception of the index mask option (-m) set to 22 positions with no mismatches and an index hash width (-w) of 16bp. Resulting SAM output files were converted to binary format (BAM), sorted, indexed and a mpileup BCF file produced using the SAMTOOLS library (Li et al. 2009). Variant calling and consensus sequences were derived using the mpileup2cns script as part of the VARSCAN package with –min-coverage set to 4.

### De novo assembly and Mitochondrial COI

Owing to its much higher level of variation, the mitochondrial COI gene was identified from the TRINITY *de-novo* assembly. Candidate COI contigs were identified and aligned (to a length of 1431 sites) using a custom script incorporating BLAT matching by translated amino acid, and coverage assessed by re-mapping sample reads at high stringency (minimum identity 0.97). In conjunction with coverage, the diversity of COI was then phylogenetically assessed against a large database of legacy and barcode COI sequences (Ratnasingham and Hebert 2007) to eliminate miss-indexing, contaminants and pseudogenes.

### Post-hoc and phylogenetic analyses

Summary statistics on super-reference assembly and use, and on target coverage and site states, were recorded for all exon-capture samples. Detailed *post –hoc* assessment of individual exons for coverage, changes in exon boundaries and patterns of polymorphic sites used a subset of well-sequenced samples spanning ophiuroid phylogenetic diversity and distance from SR. As our target is a modified version of the original genes with sections deleted, only a proportion of our exon boundaries are directly comparable with those of *Strongylocentrotus.* Within this scope, changes in exon target boundaries were measured by assessing the concentration of mapped read match ends and by visual inspection of BFAST BAM files in IGV 2.3.23 (Robinson et al. 2011). An exon boundary change was defined as a boundary changing position by at least a quarter of the length of the exon. By using a SR made of concatenated gene exons, intron loss can be detected when reads map across nominal exon boundaries. Exons confounded by closely related paralogs, such as pseudogenes, were assessed by tracking mapped exon polymorphism levels and ASR sites with different competing contigs, and comparing observed polymorphism to coalescent expectations among independent loci and samples (Hudson 1991). To provide a consistent measure of distance between sample and reference, we use a subset of 34 exons spanning 24kb that were reliably recovered and give distances only slightly less (~85%) than the whole target.

After excluding poorly-captured and otherwise dubious exons, data matrices including the 52 original transcriptome taxa were subjected to phylogenetic analysis. Owing to the size of these data matrices (100+ Megabytes) we used RAxML, applying a codon position partition (first-second and third positions) model and rooted according to previous transcriptome analyses (O’Hara et al. 2014b). Such exon-capture datasets present a large computational task requiring many hundreds or thousands of CPU hours. Therefore an approximate approach was investigated eschewing the full ML search, running only the RAxML GTR-CAT model fast bootstraps. An all-compatible consensus topology was then derived (via PAUP), and full GTR-gamma model ML branch lengths estimated in RAxML only for this consensus topology.

Custom scripts and exon-capture phylogenomic data will be available via TreeBase or Dryad (to be confirmed).

## SUPPLEMENTARY MATERIAL

Supplementary Table S1 is available from the authors.

## ACKNOWLEDGEMENTS

Thanks to Sadie Mills (NIWA), Marc Eléaume (MNHN) and Gustav Paulay (UF) for providing tissue samples, Kate Naughton (MV) for assembling plate 4, Melanie Mackenzie (MV) for facilitating transport of materials, Luisa Teasdale (MV) for bioinformatics advice, and Kevin Rowe (MV) for commenting on drafts of this paper. This work was supported by the Marine Biodiversity Hub, funded through the National Environmental Research Program (NERP), and administered through the Australian Government’s Department of the Environment. This paper is an output of the project ‘National maps of biodiversity and connectivity’.

## REFERENCES

Aberer AJ, Kobert K, Stamatakis A. 2014. ExaBayes: Massively Parallel Bayesian Tree Inference for the Whole-Genome Era. Mol Biol Evol 31:2553–2556.

Altmann A, Weber P, Bader D, Preuß M, Binder EB, Müller-Myhsok B. 2012. A beginners guide to SNP calling from high-throughput DNA-sequencing data. Hum Genet 131:1541–1554.

Bi K, Vanderpool D, Singhal S, Linderoth T, Moritz C, Good JM. 2012. Transcriptome-based exon capture enables highly cost-effective comparative genomic data collection at moderate evolutionary scales. BMC Bioinformatics 13:1–14.

Faircloth BC, McCormack JE, Crawford NG, Harvey MG, Brumfield RT, Glenn TC. 2012. Ultraconserved elements anchor thousands of genetic markers spanning multiple evolutionary timescales. Syst Biol 61:717–726.

Fell HB. 1963. The phylogeny of sea stars. Philos Trans R Soc London B 246:381–435.

Grabherr MG, Haas BJ, Yassour M, et al. 2011. Full-length transcriptome assembly from RNA-Seq data without a reference genome. Nat Biotechnol 29:644–652.

Hancock-Hanser BL, Frey A, Leslie MS, Dutton PH, Archer FI, Morin PA. 2013. Targeted multiplex next-generation sequencing: advances in techniques of mitochondrial and nuclear DNA sequencing for population genomics. Mol Ecol Res 13:254–268.

Hedtke SM, Morgan MJ, Cannatella DC, Hillis DM. 2013. Targeted Enrichment: Maximizing Orthologous Gene Comparisons across Deep Evolutionary Time. PLoS ONE 8:e67908.

Homer N, Merriman B, Nelson SF. 2009. BFAST: An Alignment Tool for Large Scale Genome Resequencing. PLoS ONE 4:e7767.

Hudson RR. 1991. Gene genealogies and the coalescent process. Oxf Surv Evol Biol 7:1–44.

Jarvis ED Mirarab S Aberer AJ, et al. 2014. Whole-genome analyses resolve early branches in the tree of life of modern birds. Science 346:1320–1331.

Kent WJ. 2002. BLAT - The BLAST-like alignment tool. Genome Res 12:656–664.

Knowles LL. 2009. Statistical Phylogeography. Annu Rev Ecol Evol Syst 40:593–612.

Leaché AD, Wagner P, Linkem CW, et al. 2014. A hybrid phylogenetic-phylogenomic approach for species tree estimation in African Agama lizards with applications to biogeography, character evolution, and diversification. Mol Phylogenet Evol 79:215–230.

Lemmon AR, Emme SA, Lemmon EM. 2012. Anchored Hybrid Enrichment for Massively High-Throughput Phylogenomics. Syst Biol 61:727–744.

Lemmon EM, Lemmon AR. 2013. High-Throughput Genomic Data in Systematics and Phylogenetics. Annu Rev Ecol Evol Syst 44:99–121.

Li C, Hofreiter M, Straube N, Corrigan S, Naylor GJP. 2013. Capturing protein-coding genes across highly divergent species. Biotechniques 54:321–326.

Li HH, B., Wysoker A, Fennell T, Ruan J, Homer N, Marth G, Abecasis G, Durbin R, Subgroup GPDP. 2009. The Sequence Alignment/Map format and SAMtools. Bioinformatics 25:2078–2079.

Lohse M, Bolger AM, Nagel A, Fernie AR, Lunn JE, Stitt M, Usadel B. 2012. RobiNA: a user-friendly, integrated software solution for RNA-Seq-based transcriptomics. Nucleic Acids Res 40:W622–W627.

Lynch M. 2002. Intron evolution as a population-genetic process. Proc Natl Acad Sci USA 99:6118–6123.

Mandel JR, Dikow RB, Funk VA, Masalia RR, Staton SE, Kozik A, Michelmore RW, Rieseberg LH, Burke JM. 2014. A target enrichment method for gathering phylogenetic information from hundreds of loci: An example from the Compositae. Appl Plant Sci 2:1300085.

Martynov AV. 2010. Reassessment of the classification of the Ophiuroidea (Echinodermata), based on morphological characters. I. General character evaluation and delineation of the families Ophiomyxidae and Ophiacanthidae. Zootaxa 2697:1–154.

Mortensen T. 1932. On an extraordinary ophiurid, *Ophiocanops fugiens* Koehler. With remarks on *Astrogymnotes, Ophiopteron*, and on an albino *Ophiocoma.* Papers from Dr. Th. Mortensen’s Pacific Expedition 1914-16. LX. Vidensk Medd Dansk Naturhist Foren 93:1–21, pl. 21.

O’Hara TD. 2007. Seamounts: Centres of endemism or species-richness for ophiuroids? Glob. Ecol. Biogeogr. 16:720–732.

O’Hara TD, England PR, Gunasekera R, Naughton KM. 2014a. Limited phylogeographic structure for five bathyal ophiuroids at continental scales. Deep Sea Res. I 84:18–28.

O’Hara TD, Hugall AF, Thuy B, Moussalli A. 2014b. Phylogenomic resolution of the Class Ophiuroidea unlocks a global microfossil record. Curr. Biol. 24:1874–1879.

O’Hara TD, Rowden AA, Bax NJ. 2011. A southern hemisphere bathyal fauna is distributed in latitudinal bands. Curr. Biol. 21:226–230.

OBIS. 2014. Global biodiversity indices from the Ocean Biogeographic Information System. Intergovernmental Oceanographic Commission of UNESCO. Web. http://www.iobis.org (consulted on 2015/1/21).

Parmley JL, Urrutia AO, Potrzebowski L, Kaessmann H, Hurst LD. 2007. Splicing and the Evolution of Proteins in Mammals. PLoS Biology 5:e14.

Ratnasingham S, Hebert PDN. 2007. BOLD: The Barcode of Life Data System (www.barcodinglife.org). Molecular Ecology Notes 7:355–364.

Robinson JT, Thorvaldsdóttir H, Winckler W, Guttman M, Lander ES, Getz G, Mesirov JP. 2011. Integrative Genomics Viewer. Nat Biotechnol 29:24–26.

Roy SW. 2009. Phylogenomics: Gene Duplication, Unrecognized Paralogy and Outgroup Choice. PLoS ONE 4:e4568.

Siepel A, Bejerano G, Pedersen JS, et al. 2005. Evolutionarily conserved elements in vertebrate, insect, worm, and yeast genomes. Genome Res 15:1034–1050.

Sprinkle J, Guensburg TE. 2004. Crinozoan, blastozoan, echinozoan, asterozoan, and homalozoan echinoderms. In: BD Webby, F Paris, ML Droser, IG Percival, editors. The Great Ordovician Biodiversification Event. New York: Columbia University Press. p. 266–280.

Stamatakis A. 2014. RAxML Version 8: A tool for Phylogenetic Analysis and Post-Analysis of Large Phylogenies. Bioinformatics 30:1312–1313.

Stöhr S, Conand C, Boissin E. 2008. Brittle stars (Echinodermata: Ophiuroidea) from La Réunion and the systematic position of *Ophiocanops* Koehler, 1922. Zool. J. Linn. Soc. 153:545–460.

Stöhr S, O’Hara TD, Thuy B. 2012. Global diversity of brittle stars (Echinodermata: Ophiuroidea). PLoS ONE 7:e31940.

Swofford DL. 2003. PAUP*. Phylogenetic Analysis using Parsimony (* and other methods). Version 4. Sunderland, Massachusetts: Sinauer Associates

Thuy B, Stöhr S. 2011. Lateral arm plate morphology in brittle stars (Echinodermata: Ophiuroidea): new perspectives for ophiuroid micropalaeontology and classification. Zootaxa 3013:1–47.

Tilston-Smith B, Harvey MG, Faircloth BC, Glenn TC, Brumfield RT. 2014. Target Capture and Massively Parallel Sequencing of Ultraconserved Elements for Comparative Studies at Shallow Evolutionary Time Scales. Syst Biol 63:83–95.

WoRMS Editorial Board. 2015. World Register of Marine Species. Available from http://www.marinespecies.org at VLIZ. Accessed 2015-01–.

Zhang J. 2003. Evolution by gene duplication: an update. Trends Ecol. Evol. 18:292–298.

